# Cell tip growth underlies injury response of marine macroalgae

**DOI:** 10.1101/2021.08.28.458046

**Authors:** Maki Shirae-Kurabayashi, Tomoya Edzuka, Masahiro Suzuki, Gohta Goshima

## Abstract

Regeneration is a widely observed phenomenon by which the integrity of an organism is recovered after damage. So far, studies on the molecular and cellular mechanisms of regeneration have been limited to a handful of model multicellular organisms. Here, we systematically surveyed the regeneration ability of marine macroalgae (Rhodophyta, Phaeophyceae, Chlorophyta) after thallus severing and applied live cell microscopy on them to uncover the cellular response to the damage. We observed three types of responses – budding, rhizoid formation and/or sporulation – in 25 species among 66 examined, demonstrating the high potential of regeneration of macroalgae. In contrast, callus formation, which often accompanies plant regeneration, was never observed. We monitored the cellular and nuclear dynamics during cell repair or rhizoid formation of four phylogenetically diverged Rhodophyta and Chlorophyta species (*Colaconema* sp., *Dasya sessilis, Cladophora albida, Codium fragile*). We observed tip growth of the cells near the damaged site as a common response, despite the difference in the number of nuclei and cells across species. Nuclear translocation follows tip growth, enabling overall uniform distribution of multinuclei (*Dasya sessilis, Cladophora albida, Codium fragile*) or central positioning of the mononucleus (*Colaconema* sp.). In contrast, the control of cell cycle events, such as nuclear division and septation, varied in these species. In *Dasya sessilis*, the division of multinuclei was synchronised, whereas it was not the case in *Cladophora albida*. Septation was tightly coupled with nuclear division in *Colaconema* and *Dasya* but not in others. These observations show that marine macroalgae utilise a variety of regeneration pathways, with some common features. This study also provides a novel methodology of live cell biology in macroalgae, offering a foundation for the future of this under-studied taxon.

## Introduction

Regeneration is widely observed in multicellular organisms; injured tissue (*e*.*g*. the human liver and lizard’s tail) or, in some cases, the whole body of an organism (*e*.*g*. planaria and moss) is eventually recovered through cell proliferation and differentiation (Duncan and Sanchez Alvarado, 2019; Murawala et al., 2012). Marine organisms often incur injuries, typically caused by storms or predators. As a result, they have certain response mechanisms to this damage. However, the regeneration process has hardly been described at a cellular level for marine creatures, and the underlying mechanisms remain poorly understood.

Marine macroalgae, or seaweeds, are important players in marine ecology (Graham et al., 2008). Three types of seaweeds, namely, red (Rhodophyta), brown (Phaeophyceae), and green (Chlorophyta) macroalgae, are widely distributed, particularly in coastal areas. Phylogenetically, red and green algae and land plants constitute a single clade, whereas brown algae are more distinct (Adl et al., 2019). Their ability to photosynthesise makes them critical CO2 consumers and oxygen producers. They also serve as food sources and habitats for benthic and nektonic animals, and the interaction with animals raises the risk of injury. A high potential for regeneration has been described in macroalgal species. When a filament of *Griffithsia pacifica* Kylin was severed, regrowth of the cell next to the damaged cell was observed (Waaland and Cleland, 1974). In the red alga *Gracilariopsis chorda* (Holmes) Ohmi, experimentally severed thalli were returned to the ocean, and vegetative regeneration was subsequently observed (Kim et al., 2005). Furthermore, in the brown alga *Dictyota dichotoma* (Hudson) J.V. Lamouroux, budding and rhizoid formation from a severed surface has been reported in juvenile thalli (Tanaka et al., 2017). *In vitro* regeneration assays have been also conducted, which involved artificially prepared protoplasts or the callus (Baweja et al., 2009; Mine et al., 2008). Studies show that protoplasts of the green algae *Ulva* (sea lettuce) and *Bryopsis* regenerate to form the thallus (Fujimura et al., 1989; Tatewaki and Nagata, 1970). Explants of gracilarialean red algae generate calli, which are stimulated by plant growth hormones (Gusev et al., 1987; Yokoya et al., 2004). However, the use of different assays in different studies makes it difficult to deduce whether there is a common mode of regeneration in macroalgae. Furthermore, how cells initially respond to injury remains largely unclear at both the cellular and intracellular level, as high-resolution time-lapse imaging of the cellular and intracellular components has scarcely been applied.

In this study, we conducted a systematic survey of the response to injury (which we introduced by thallus severing) of 66 locally obtained macroalgal species. Thereafter, we analysed cell growth, cell division, and nuclear dynamics. Nuclear dynamics are of particular interest in macroalgae, as many species have multiple nuclei within one cell. Our survey identified three major macroscopic responses to thallus severing, namely, bud, rhizoid, and spore formation. At the cellular level, instant cell tip growth, accompanied by nuclear translocation, was commonly observed upon injury in four red/green algal species.

## Materials and methods

### Macroalgae collection

Macroalgae were predominantly collected from the intertidal zone of the seashore, in front of the laboratory (Lat. 34°29’8” N, Long. 136°52’32″ E, Fig. 1A–C) on the 25^th^ and 26^th^ of March 2020, and on the 30^th^ of March and 1^st^ of April 2021. Samples were also collected from the outside tank of the laboratory, as well as from the ropes of the floating pier (Fig. 1C–F). The macroalgae were kept in natural seawater at 15 °C until they were severed.

**Figure 1.**
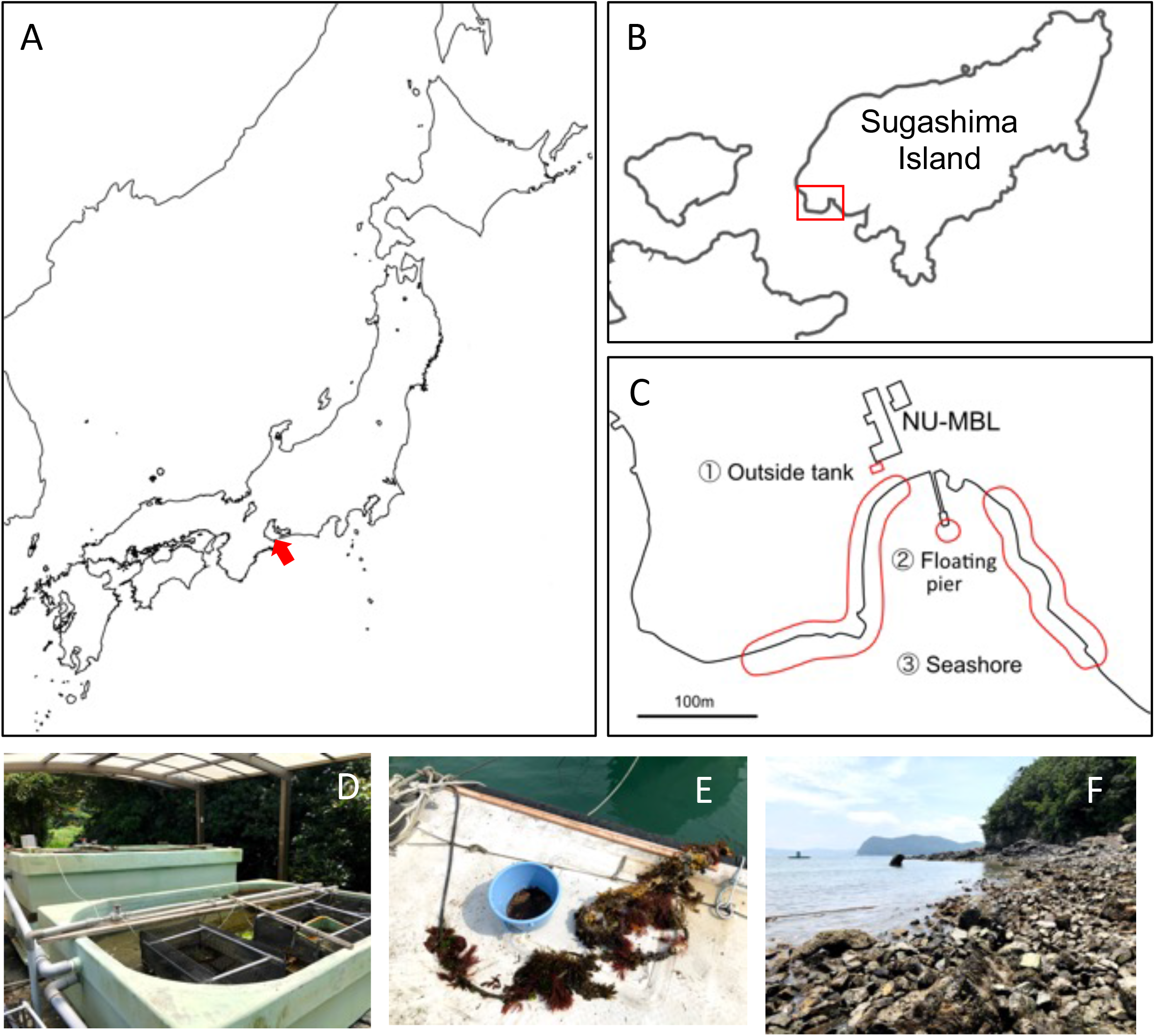
Macroalgae collection sites. (A–C) Location of the Sugashima Marine Biological Laboratory (NU-MBL) and sample collection sites (indicated by red marks). (D–F) The three sites of macroalgae collection. Outdoor tank at NU-MBL which has a continuous flow of unfiltered seawater (D), the underwater rope at the pier was a source of macroalgae (E), and the intertidal zone in front of the NU-MBL (F).

### Identification of macroalgae

Collected macroalgae were examined to determine their probable genera and/or species; monographic publications and floristic studies were used to confirm the morphology and identify the species (Takamine and Yamada, 1950; Yoshida, 1998). The nomenclature used in this study follows Algaebase (Guiry and Guiry, 2021). DNA barcode sequencing was performed for some specimens to confirm and/or correct the morphological identification. The plastid *rbc*L, mitochondrial cox1, nuclear 18S rDNA, and/or rDNA ITS1-5.8S-ITS2 region were amplified and sequenced. Genomic DNA was extracted using the DNeasy Plant Mini Kit (Qiagen, Tokyo, Japan) from field-collected specimens dried by silica gel. Total DNA was used as the template for polymerase chain reactions (PCR) in which the *rbc*L, cox1, 18S rDNA, and/or rDNA ITS1-5.8-ITS2 loci were amplified using a KOD-ONE kit (Toyobo Co. Ltd., Osaka, Japan) and a 2720 PCR Thermal Cycler (Applied Biosystems, Foster City, CA, USA). The primers used for PCR amplification are shown in Table 1. The temperature cycling protocol was as follows: 2 min at 94 °C for an initial denaturation step, 35 cycles of 15 s denaturation at 94 °C, 30 s primer annealing at 46 °C, and 1 min extension at 68 °C, with a final 7 min extension at 72 °C, followed by a hold at 4 °C. The amplified DNA fragments were purified by ethanol precipitation and sequenced by eurofins genomics (Luxembourg). Reverse and direct chromatograms were assembled using ClustalW software.

**Table 1:**
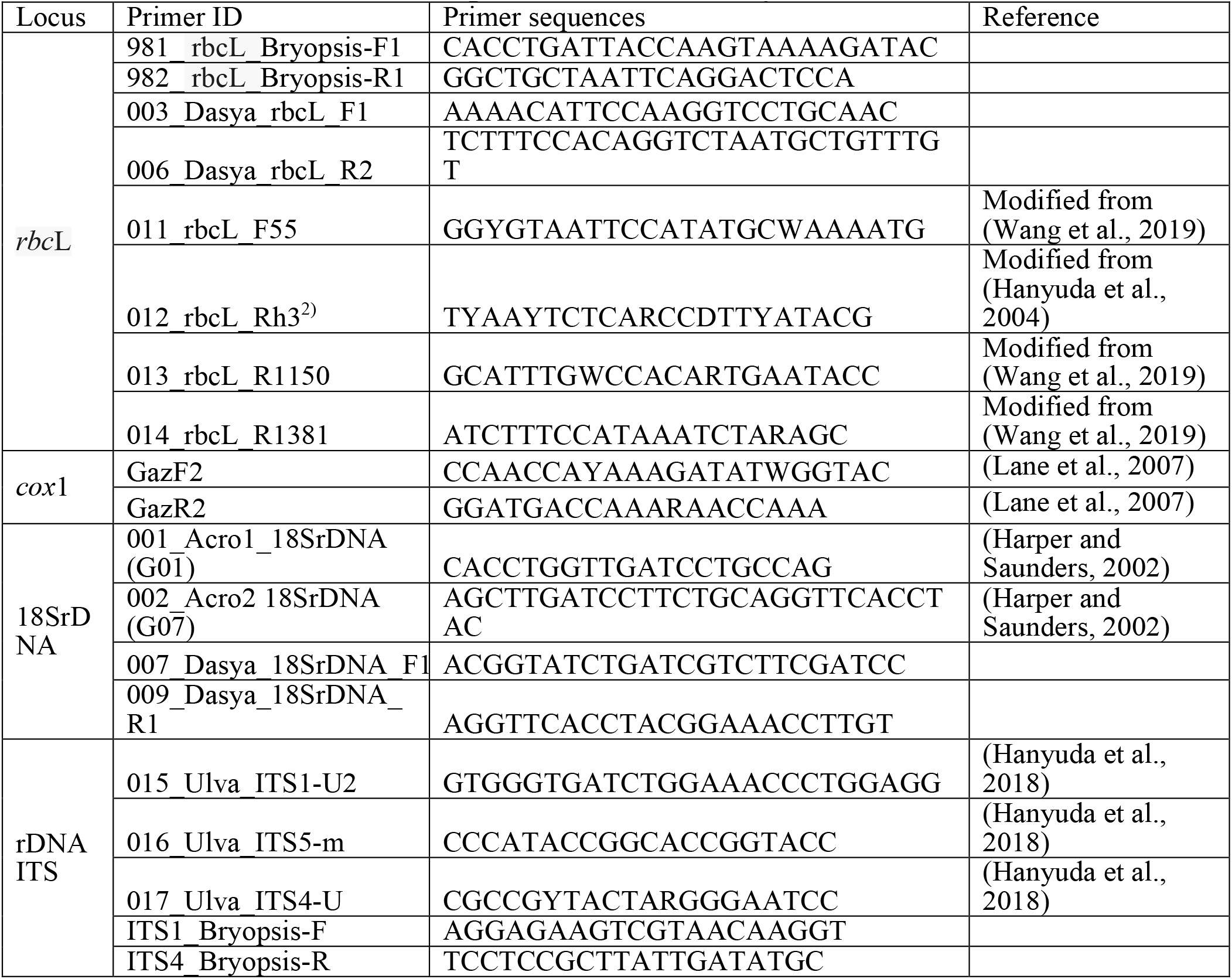
Primers used for PCR amplification in this study.

### Severing experiments

The thalli of each species were washed once with filtered seawater and severed in approximately 5 mm (× 1–5 mm) fragments. The fragments were immersed in modified seawater (MSW), which was prepared using the following method: ocean surface water was filtered using a 0.45-µm Millipore Stericup, autoclaved, and supplied with GeO2 (final concentration, 1 mg/L) and Provasoli’s enriched seawater (PES) or Daigo’s IMK medium for long-term culture (25 °C, always expose light 1200 lux.). We checked the severed site of the algae with a stereomicroscope every day for a maximum of 21 days. For frozen sections of *Gracilariopsis chorda*, the served thalli were fixed with 4% paraformaldehyde in PHEM buffer (60 mM Pipes, 25 mM Hepes, 10 mM EGTA, 2 mM MgCl2, pH 6.9) and frozen with OCT medium at -80 °C for 1 h. Sections were prepared using CM3000 Cryostat (Lica biosystems, Wetzlar, Germany).

### Imaging

Stereomicroscopic images were acquired using an ILCE-QX1 camera (Sony, Tokyo, Japan) attached to SMZ800N microscope (Nikon). A Plan 1× lens was used. Section imaging was acquired with an ILCE-QX1 camera (Sony) attached to ECLIPSE E200 (Nikon). For time-lapse microscopy, thalli were severed into < 2 mm pieces in the MSW and treated with Hoechst 33342 at 10 µg/mL (final) for DNA and cell wall staining. After 30 min incubation with the Hoechst dye, the thalli were injected into the microfluidic device which has previously been used for moss imaging (Kozgunova and Goshima, 2019). In brief, a polydimethylsiloxane (PDMS) device was attached to a glass-bottom dish (dish diameter 35 mm; glass diameter 27 mm; glass thickness 0.16–0.19 mm), into which severed thalli were injected by needle (Fig. S1). The height of the device was 15 µm. Hoechst imaging was performed at 23 °C with a Nikon inverted microscope (Ti2) attached to a 40× lens (0.95 NA), a 405-nm laser, a CSU-10 spinning-disc confocal unit, and a CMOS camera (Zyla, Andor). Depending on the species under observation, sequential image acquisition time intervals varied between 10 and 30 min. The microscopes were controlled using NIS Elements software (Nikon). The data were analysed using FIJI.

## Results

### Budding, rhizoid formation, and sporulation are three major injury responses

Among the macroalgal thalli collected in front of our marine laboratory (Fig. 1), we identified 54 species through visual inspection (26 red, 21 brown, and 7 green algae). In addition, we collected 14 species that could not be morphologically identified (Table 2). Consequently, DNA barcode sequences were determined for these unidentified species and compared with those registered in the database (Table S1). Two species died before the experiment, so we conducted serving experiments on 66 species.

**Table 2.**
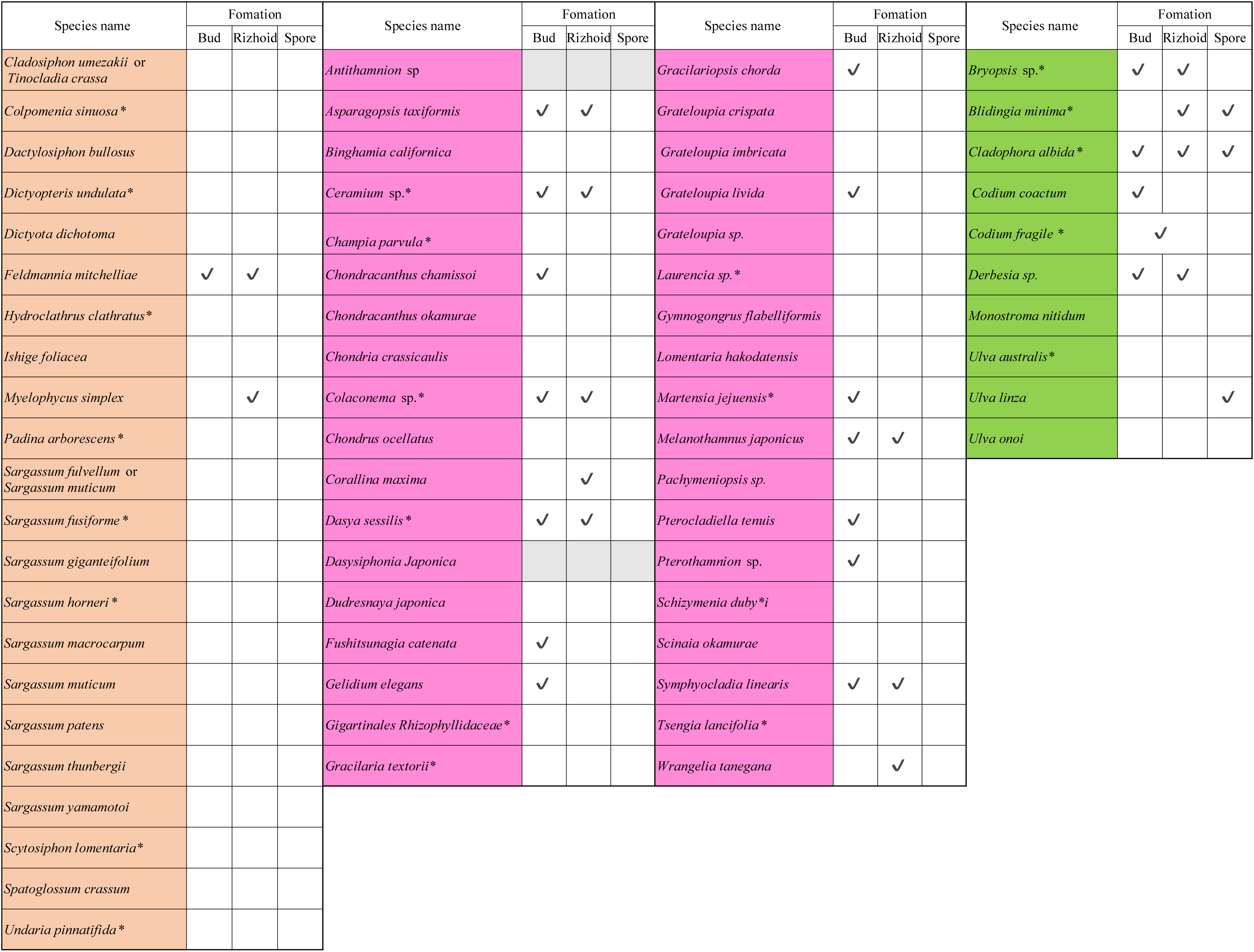
Summary of the response to thallus severing. Grey: untested.

Results from the 21 day-follow up of the severed sites in the algae are summarised in Fig. 2A. We observed immediate death or no specific response for 41 species, whereas the other 25 species showed three major responses: rhizoid formation (yellow arrowheads in Fig. 2B, 2E, and 2F), budding (white arrowheads in Fig. 2C–F), and sporulation (Fig. 2G). Two or three of these responses were simultaneously observed in 11 species (Fig. 2E, F). Even though many specimens were severed in multiple locations, we did not find any site-dependent responses. There was a clear tendency in the response type for each algal lineage. The majority of the brown algae showed no response (2/22 formed buds and/or rhizoids) (Fig. 2A). In contrast, 47% (16) of the red algae specimens showed budding and/or rhizoid formation. Sporulation was only observed in green algae (3/10 showed sporulation). In all cases, buds and rhizoids were formed at the apical and basal regions, respectively, indicating that the apicobasal polarity was maintained after being severed (Fig. 2F). Depending on the species, buds were observed on the inner part of the severed surface (Fig. 2C, S2) or at its edge (Fig. 2D). During regeneration of buds, we could not observe callus-like cell masses. Rhizoid formation was observed at the basal severed site (Fig. 2B, 2F). Thus, our survey revealed that macroalgae have a high regeneration potential, that is limited to three patterns, budding, rhizoid formation, and sporulation.

**Figure 2.**
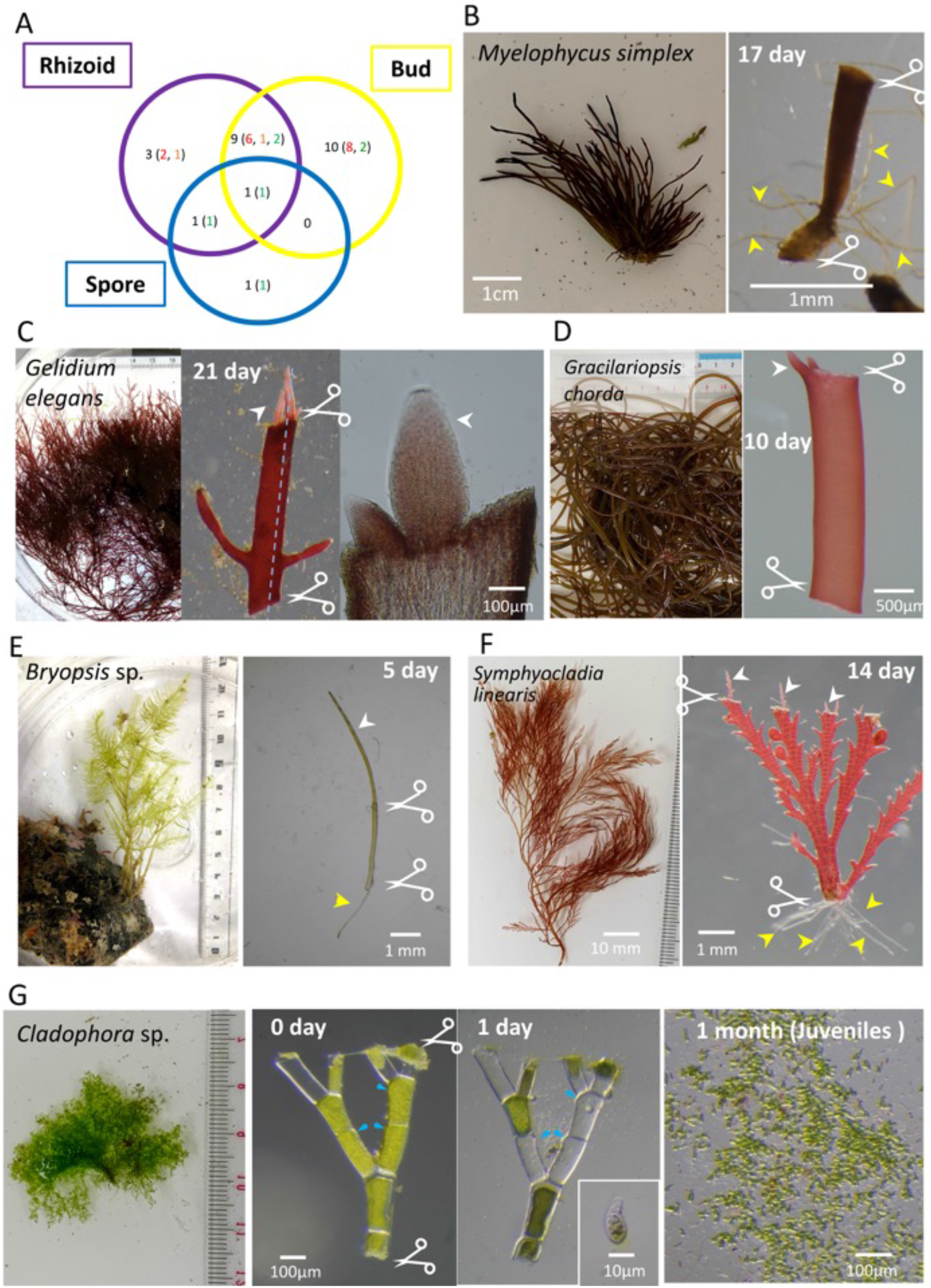
Budding, rhizoid formation, and sporulation are the major responses to injury. (A) Number of macroalgal species that showed the indicated response. The numbers of red, brown, and green algae are indicated by their colours. (B) Rhizoid formation at the basal cut site in *Myelophycus simplex*. (C) Budding at the cut site of *Gelidium elegans*. The rightmost panel is a section image of the blue dotted line in the second panel. Buds appeared in the inner part of the cut surface. (D) Buds appeared at the edge of the cut surface in *Gracilariopsis chorda*. (E) Bud and rhizoid formation after cutting of *Bryopsis* sp. (F) Bud and rhizoid formation after cutting of *Symphyocladia linearis*. (G) Sporulation in *Cladophora albida*. Within 24 h after injury, spores were formed and released from the thalli. White arrowhead, bud; yellow arrowhead, rhizoid; blue arrowhead, pore through which spores are released.

### Regeneration in uninuclear *Colaconema* sp. is characterised by cell tip growth and nuclear division, followed by septation

We were interested in the cellular and intracellular responses to injury and studied four species in more detail, namely, *Colaconema* sp. (red), *Dasya sessilis* Yamada (red), *Cladophora albida* (Nees) Kützing (green), and *Codium fragile* (Suringar) Hariot (green). The specimens were obtained during the springtime of two successive years and were kept in the laboratory tank for >1 year. This meant that experiments were conducted repetitively.

*Colaconema* sp. is a filamentous red alga obtained from the surface of *Codium* (Fig. 3A). However, it grew well in seawater without *Codium* in the laboratory. Fig. 3B shows rhizoid formation in the basal region after severing the thallus.

**Figure 3.**
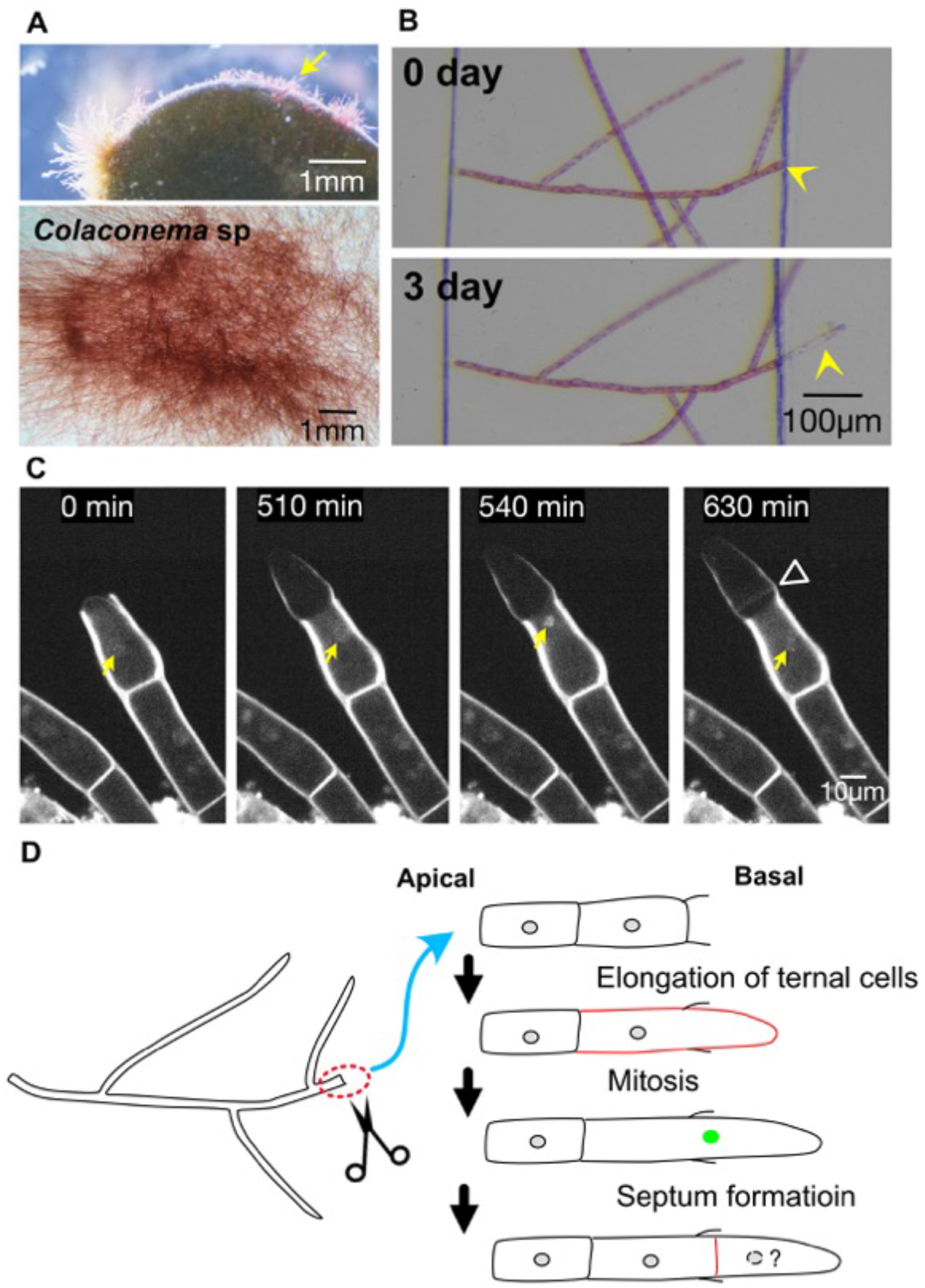
Tip growth, nuclear division, and septation in *Colaconema* sp. (A) *Colaconema* sp. was isolated from the surface of *Codium fragile* (top). *Colaconema* sp. was successfully cultured in the absence of *C. fragile* in the laboratory (bottom). (B) Rhizoid formation upon severing. (C) Time-lapse imaging of the nuclei (yellow arrows) of a regenerating cell (rhizoid). The cell next to the injured cell underwent a nuclear division followed by septation (arrowhead) during tip growth. Mitotic chromosome condensation was observed at 540 min, and sister chromatid segregation was complete by 630 min (a sister nucleus is out of focus). (D) Schematic presentation of the injury response in *Colaconema* sp. Mitotic nuclei are indicated in green.

To investigate the cellular and intracellular responses during rhizoid formation upon injury, we stained the nucleus with Hoechst 33342 and performed time-lapse imaging at 30 min intervals in a 15-µm height microfluidic chamber (Fig. S1). The specimen was kept close to the glass surface, making higher-resolution imaging applicable (Fig. 3C, Video 1). Consistent with the DAPI staining of fixed *Colaconema caespitosum* (J. Agardh) Jackelman, Stegenga & J.J. Bolton (Garbary and Zuchang, 2006), we observed a single nucleus in each cell. The cell next to the injured site started to regrow. Concomitantly, the nucleus was translocated apically, and remained in a position that was roughly central in the cell. At 540 min, chromosomes were condensed, followed by sister chromatid separation and segregation. Septation occurred immediately after chromosome segregation (Fig. 3C, arrowhead). Fig. 3D schematically describes the injury response.

### Regeneration of multinuclear *Dasya sessilis* starts with cell death, followed by neighbour growth, synchronous nuclear division, and septation

The red alga *D. sessilis* develops a cylindrical erect axis, with branches gradually becoming shorter towards the top. Each branch is covered with dense adventitious pseudo-laterals composed of multinucleated cells (Fig. 4A). Upon severing of the thallus, new cells were observed in the apical and basal regions and were morphologically indistinguishable. However, only the cells on the apical side formed branches on day 5, indicating that buds and rhizoids formed in the apical and basal regions, respectively (Fig. 4B).

**Figure 4.**
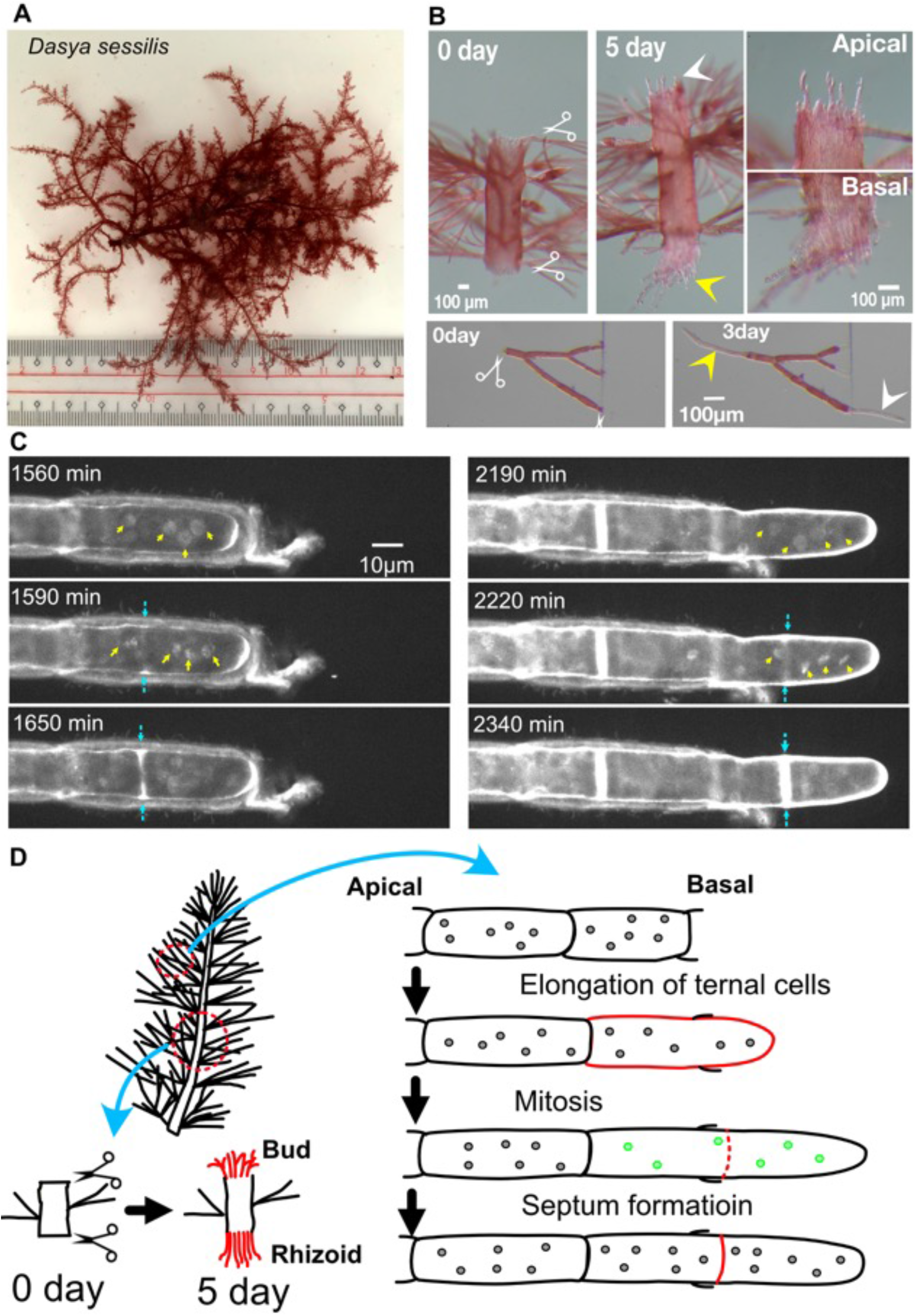
Tip growth, synchronised multinuclear division, and septation in *Dasya sessilis*. (A) *Dasya sessilis*. (B) Formation of buds (white arrowheads) and rhizoids (yellow arrowheads) at the apical and basal cut sites, respectively. (C) Time-lapse imaging of the nuclei of a regenerating cell (rhizoid). The cell next to the injured cell underwent two rounds of nuclear divisions and septation during tip growth. Yellow and blue arrows indicate mitotic chromosome condensation and septation, respectively. (D) Schematic presentation of the injury response in *Dasya sessilis*. Mitotic nuclei are indicated in green.

To investigate the cellular and intracellular responses to severing, particularly regarding how rhizoids are formed, we stained the nuclei with Hoechst 33342 and performed time-lapse imaging at 30 min intervals in the microfluidic chamber (Fig. 4C, Video 2). In comparison to the clarity obtained with cell wall staining, nuclear staining was not very clear in *D. sessilis*. Nevertheless, consistent with a previous description (Graham et al., 2008), we observed multiple nuclei in each cell (Fig. 4C, arrows). The severed cell immediately died, suggesting that the cell repair mechanisms observed in some coenocytes (Mine et al., 2008) do not operate in *D. sessilis*. However, the cell adjacent to the injured site started to show tip growth, and concomitantly the nuclei moved in an apical direction (Fig. 4C). At 1590 and 2220 min, the chromosomes of all four nuclei in the focal plane were condensed (Fig. 4C). We were able to trace the entire process in three cells, all of which showed synchronous mitotic chromosome condensation. Thus, mitosis is likely to be synchronised in *D. sessilis*. The precise timing of chromosome segregation was not visible at our time resolution; nevertheless, we were able to determine that it occurred within 30 min after mitotic entry, based on the disappearance of condensed chromosomes. Septation occurred within 1 h of chromosome segregation (Fig. 4C, blue arrows). Despite the multiple nuclear segregation events that occurred at one time, the septum was formed only at one site, preserving the multinuclear state of the cell after septation (Fig. 4D).

### Regeneration of multinucleate *Cladophora albida* is characterised by simultaneous tip growth of apical and subapical cells, followed by partially synchronous multinuclear division

The green algae *Cladophora albida* belongs to the family Cladophoraceae. As typical to this family, it develops filamentous, monosiphonous thalli with branches composed of multinucleate cells. Upon severing, all three major responses were observed (Figs. 2E — sporulation— and 5A —bud and rhizoid formation). As previously seen in *Cl. glomerata* (Linnaeus) Kützing using DAPI staining (Zulkifly et al., 2013), we observed multiple nuclei in a cell (Fig. 5B–D). The cell next to the severed site started to regrow rapidly (1. 9 ± 0.5 µm/min, n = 14). In addition, adjacent cells further away from the injured site regrew at a slightly slower rate (0.9 ± 0.3 µm/min, n = 9), which was neither observed in *Colaconema* sp. nor *D. sessilis*. In most cases, the cells next to the severed site elongated without nuclear division, and the vacuoles developed rapidly, occupying the cytoplasm in the elongated cells (Fig. 5B, Video 3). In two cases, however, we observed many nuclear division events in the elongated cells (Fig. 5C, coloured arrows, Video 4). We found that not all nuclear divisions were entirely synchronised. While synchronous nuclear divisions were observed for some nuclei, others divided at different time points. This is different from that observed in *D. sessilis*, in which all nuclear divisions within a cell were synchronised. Septation occurred infrequently (Fig. 5C, white arrowheads).

**Figure 5.**
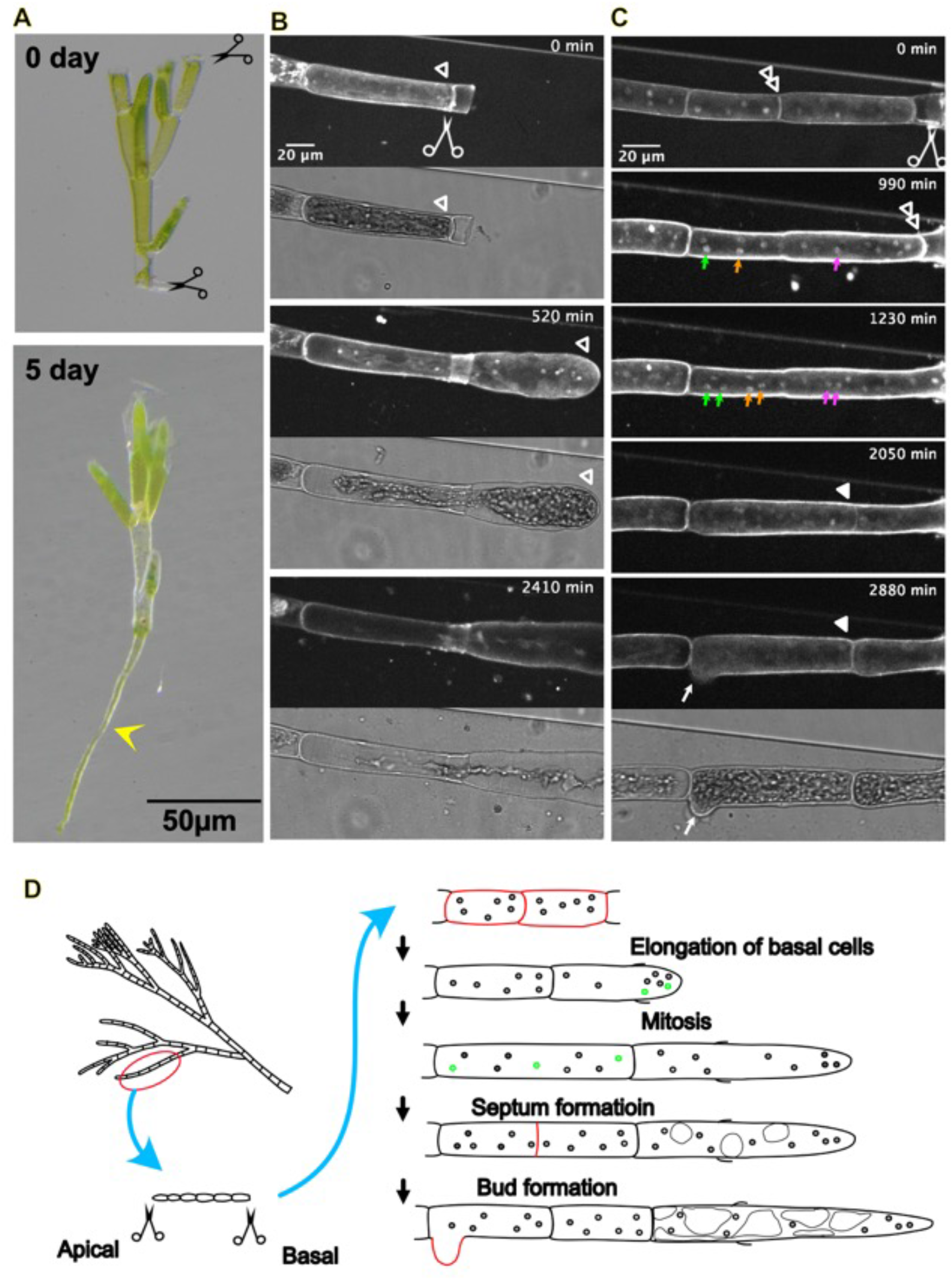
Tip growth, unsynchronised multinuclear division, and septation in *Cladosphora albida*. (A) Formation of rhizoids (yellow arrowhead) at the basal cut site. (B, C) Time-lapse imaging of the nuclei of a regenerating cell (rhizoid). The cell next to the injured cell showed tip growth (tips are indicated by open white arrows). (B) In the majority of cases, nuclear division was undetectable, whereas the vacuole developed and occupied the cytoplasm. (C) In other cases, the growing cell underwent nuclear division, not in a synchronous manner, followed by single septation during tip growth. Branching is also observed at the end (white arrow). Green, yellow, and magenta arrows indicate mitotic chromosome condensation, whereas septation is indicated by filled white arrowheads. (D) Schematic presentation of the injury response in *Cladosphora* sp. Mitotic nuclei are indicated in green.

### Regeneration of *Codium fragile* involves cell repair and new tip growth, followed by nuclear translocation without division

The green algae *Codium fragile* belongs to family Codiaceae, In *C. fragile*, the thick branches are composed of closely packed utricles, which consist of small cylindrical club-shaped structures (Fig. 6A, top right). The structure is characterised by an ellipsoidal portion and two associated medullary filaments (Fig. 6A, bottom left). The cytoplasm is continuous, with only an incomplete septum between the filaments and the ellipsoidal portion (Fig. 6A, blue arrow) (Nanba et al., 2002; Nanba et al., 2000). The structure could be maintained in a petri dish with MSW for less than two weeks; thereafter, the ellipsoidal portion lost the plastids while medullary filaments started to grow (Fig. 6A, bottom right). The filament kept growing for more than a year in the petri dish.

**Figure 6.**
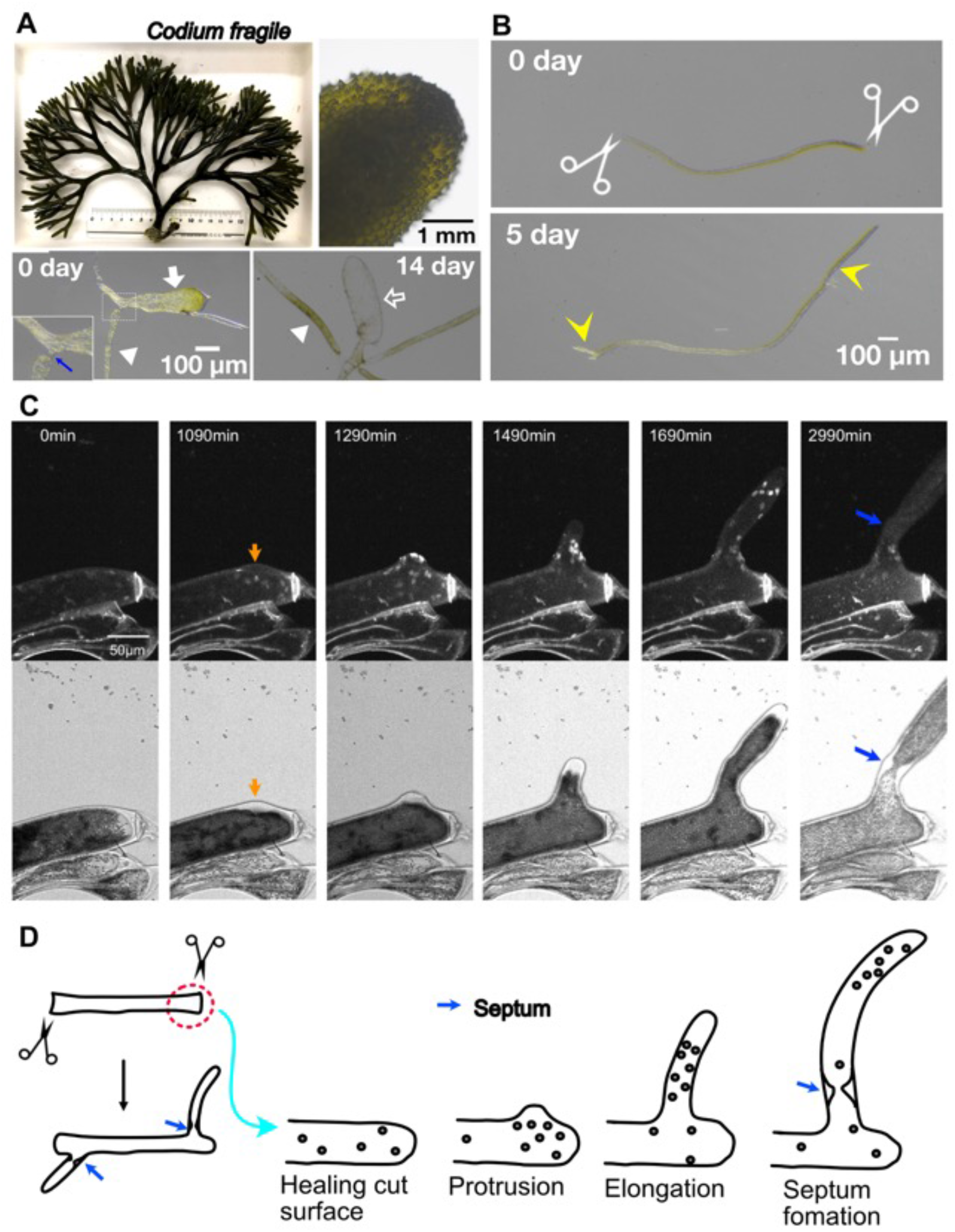
Cell repair, protrusion, tip growth, and nuclear migration in *Codium fragile*. (A) *Codium fragile*. Close-up of the thalli that are composed of closely packed utricles (top right). A single utricle (bottom left, white arrow) with an incomplete septum (blue arrow) and tubular branches (white arrowhead). Fourteen days after isolation, the utricles withered (white open arrow), and the tubular branches grew to form filamentous thalli. (B) Formation of rhizoids (left arrow) and new branches (right arrow) after severing. (C) Time-lapse imaging of nuclei (dyed with Hoechst 33342) (top) and plastids that are visualized by transmission light microscopy (bottom). The cut-repaired site is located on the right (indicated by a strong, linear Hoechst signal). The orange arrow indicates the emergence of a bud. Blue arrows indicate incomplete septation. (D) Schematic presentation of the injury response of *C. fragile*.

We conducted a severing experiment on isolated filamentous thalli. Upon severing of a filament at two sites, we observed a unique response, in which the severed site was immediately (< 30 min) repaired or enclosed and growth was observed at both ends (Fig. 6B). Hoechst staining confirmed the multinucleate nature of the filaments (Fig. 6C, D, Video 5). After 20 h, protrusion and subsequent tip growth were observed adjacent to the cut site (Fig. 6C, orange arrow). In contrast to *D. sessilis* and *Cl. albida*, mitotic chromosome segregation or chromatid segregation was not observed in the severed area. Interphase nuclei were translocated into the newly formed protrusions without mitosis (1490–1690 min). Once the protrusion grew to a certain length, incomplete septation was observed, through which large organelles such as the nucleus could no longer pass (Fig. 6C, blue arrow).

## Discussion

### Macroalgal response to injury

To systematically study the injury response of macroalgae, we conducted a single thallus severing assay for 66 macroalgal species and observed the response at the macroscopic and, for some species, cellular and intercellular levels. Three responses were observed, namely, budding, rhizoid formation, and sporulation. The dominant response depended on the species. Nevertheless, all these responses would be reasonable for macroalgal survival in the natural environment. Rhizoid formation, which often occurred within 3 days, would allow the severed thallus to quickly re-attach to the substrate, while budding efficiently restored the cell types that had been lost by severing. Sporulation occurred within 1 day (*Cl. albida*) or 14–21 days (*Ulva linza* Linnaeus), enabling the transportation and development of a new thallus. It should be noted, however, that the ‘negative’ results observed in the assay (either death or no specific response) in 41 different samples were not conclusive; we hypothesize that the response-ability might be endowed to every species but could not be revealed in our specific experimental conditions (*e*.*g*. lack of water flow and the irregular lighting). Furthermore, although sporulation was observed only in green algae in this study, the same response has been reported upon wounding or heat stress in Rhodophyta (Suda and Mikami, 2020), and this response may be dependent on the age of the thalli. Besides, rhizoid and bud formation have been reported to occur in the juveniles of some Phaeophyceae species, such as *Sargassum* and *Dictyota* (Tanaka et al., 2017; Uji et al., 2015). However, we failed to observe the response for most Phaeophyceae, including *Sargassum*. Thus, the frequency of regenerating species may be underestimated, and this study reinforces the idea that macroalgae have a high regeneration rate (Baweja et al., 2009; Huang and Fujita, 1997).

An unexpected finding in this study was that we were unable to observe the formation of callus, the cell mass containing undifferentiated cells. The buds formed at the severed site in Rhodophyta *Gelidium elegans* Kützing and *Gr. chorda* were organised in form, thus suggesting the absence of callus formation. Callus formation is the best-known wound-healing response of land plants and has also been reported in many macroalgal species (Baweja et al., 2009; Ikeuchi et al., 2016; Perez-Garcia and Moreno-Risueno, 2018). However, many studies on callus formation in macroalgae are based on *in vitro* tissue cultures, often supplied with extrinsic compounds such as high doses of plant hormones (Baweja et al., 2009). Our observation of the cellular response of four macroalgal species also supports the notion that the callus is not a general intermediate during macroalgal regeneration after injury.

### Injury response at the cellular and intracellular level

To the best of our knowledge, this study represents the first report of the nuclear dynamics of macroalgae, based on live cell imaging. Generally, fluorescence imaging is challenging for photosynthetic organisms because of the high autofluorescence derived from plastids. Macroalgae are not an exception, and we detected very high signals when the samples were imaged with 560 nm and 640 nm lasers, and to a lesser extent, with a 488 nm laser; this last laser is routinely used for GFP imaging. However, autofluorescence was negligible when cells were illuminated with a 405 nm laser, allowing nuclear staining by Hoechst 33342, which is permeable to many, if not all, cell types. Hoechst staining also stained the cell wall, serving as a clear indicator of the cell growth direction and the timing of septation. One caveat of Hoechst imaging is that the low-wavelength laser confers toxicity to cells. In our case, this was minimised by lowering the laser power and exposure time as much as possible and allowing sufficient off-time between image acquisitions. The use of microfluidic devices also made a critical contribution to live imaging, as the nuclear image could be obtained by single focal plane imaging (*i*.*e*. minimal laser exposure). We believe that the intact cellular state was maintained during our imaging, as evidenced by the occurrence of mitotic cell division (in three species) and nuclear translocation (in *Codium*). We propose that the current method can be applicable to the nuclear imaging of many macroalgal species.

In three species, the injured cells were abandoned irrespective of the number of intact nuclei retained within them, and the neighbouring cells started the reaction. The injury might be recognised by the neighbouring cells, the cytoplasm of which might be connected through pit plugs, a structure formed between adjoining cells during an incomplete septation in cytokinesis in red algae (Pueschel and Cole, 1982). *Codium* was distinct in that it quickly repaired the injured site, perhaps by membrane closure. The function of the incomplete septum (‘broken wall’) in this process remains unclear. A similar response was observed for *Bryopsis*, which belongs to the same order as *Codium*. A highly viscous cytoplasm might underlie this quick repair of large injuries. The next step was tip growth of the injured or neighbouring cells. Nuclear migration then followed to maintain the relative position of the nucleus in a cell. This behaviour is similar to what has been observed in the multinucleated xanthophycean algae *Vaucheria*, where the nuclei are moved towards the branches that have been exposed to blue light (Takahashi et al., 2001). The tip growth and nuclear translocation were commonly observed in all four species. In contrast, there appeared to be a substantial variation in cell cycle control. While *Dasya* cells underwent synchronous nuclear division, this was not the case in *Cladophora*. In *Cladophora*, the nuclei that were closely located to each other tended to undergo synchronous division, whereas those in other areas did not, despite sharing the same cytoplasm. Single-nucleated *Colaconema* was similar to *Dasya*, in the sense that tip growth and nuclear motility were coupled. *Codium*, however, is extremely different, in that mitosis has been scarcely observed during regeneration. In our imaging attempts (> 10), we observed only one nuclear division among > 100 nuclei. The injury response of *Codium* may represent cellular wound healing, which does not affect cell cycle control. Finally, we found that the formation of the septum, including the incomplete septum in *Codium*, was a variable process. Interestingly, in Rhodophyta *Colaconema* and *Dasya*, this was tightly coupled with nuclear division and formed immediately after nuclear segregation. In contrast, septum formation was loosely coupled with nuclear division in (multinucleated) Chlorophyta, *Cladophola*, and *Codium*. In the multinucleated fungus *Aspergillus nidulans* G Winter, septation occurs once the tip has grown to a certain length (Wolkow et al., 1996). However, this does not appear to hold true for *Cladophora*, as we could not infer whether and when septation takes place based on cell length (Fig. 5B, C). The regulation of septum formation is an interesting topic for future research.

## Acknowledgments

We thank Noiri Oguri, Daisuke Iijima, and Gakuho Kurita (Nagoya University) for assistance with macroalgae collection and severing, Masashi Fukuoka (NU-MBL) for macroalgae maintenance, Takeaki Hanyuda and Hiroshi Kawai (Kobe University, Japan) for instructions on basic seaweed handling, Yukimasa Yamagishi (Fukuyama University), and Toyoki Iwao (Toba City Fisheries Research Center) for helping with species identification. This work was supported by a JSPS KAKENHI grant (17H06471 to G.G., 21H04162 to M.S-K., and 20K06797 to M.S.). The authors declare no conflicts of interest.

## Video legends

**Video 1: Tip growth, nuclear division, and septation in *Colaconema* sp**.

Time-lapse imaging of Hoechst 33342 using spinning-disc confocal microscopy. Arrow indicates condensed chromosomes in mitosis.

**Video 2. Tip growth, synchronised multinuclear division, and septation in *Dasya sessilis***.

Time-lapse imaging of Hoechst 33342 using spinning-disc confocal microscopy. Arrows indicate condensed chromosomes in mitosis.

**Video 3. Tip growth without nuclear division or septation in *Cladosphora albida***.

Time-lapse imaging of Hoechst 33342 using spinning-disc confocal microscopy.

**Video 4. Tip growth, unsynchronised multinuclear division, and septation in *Cladosphora albida***.

Time-lapse imaging of Hoechst 33342 using spinning-disc confocal microscopy. Arrows indicate condensed chromosomes in mitosis.

**Video 5. Tip growth, nuclear migration and incomplete septum formation in *Codium fragile***.

Time-lapse imaging of Hoechst 33342 using spinning-disc confocal microscopy.

## Tables

**Table S1**: DNA barcode sequences of several macroalgal species

**Figure S1:**
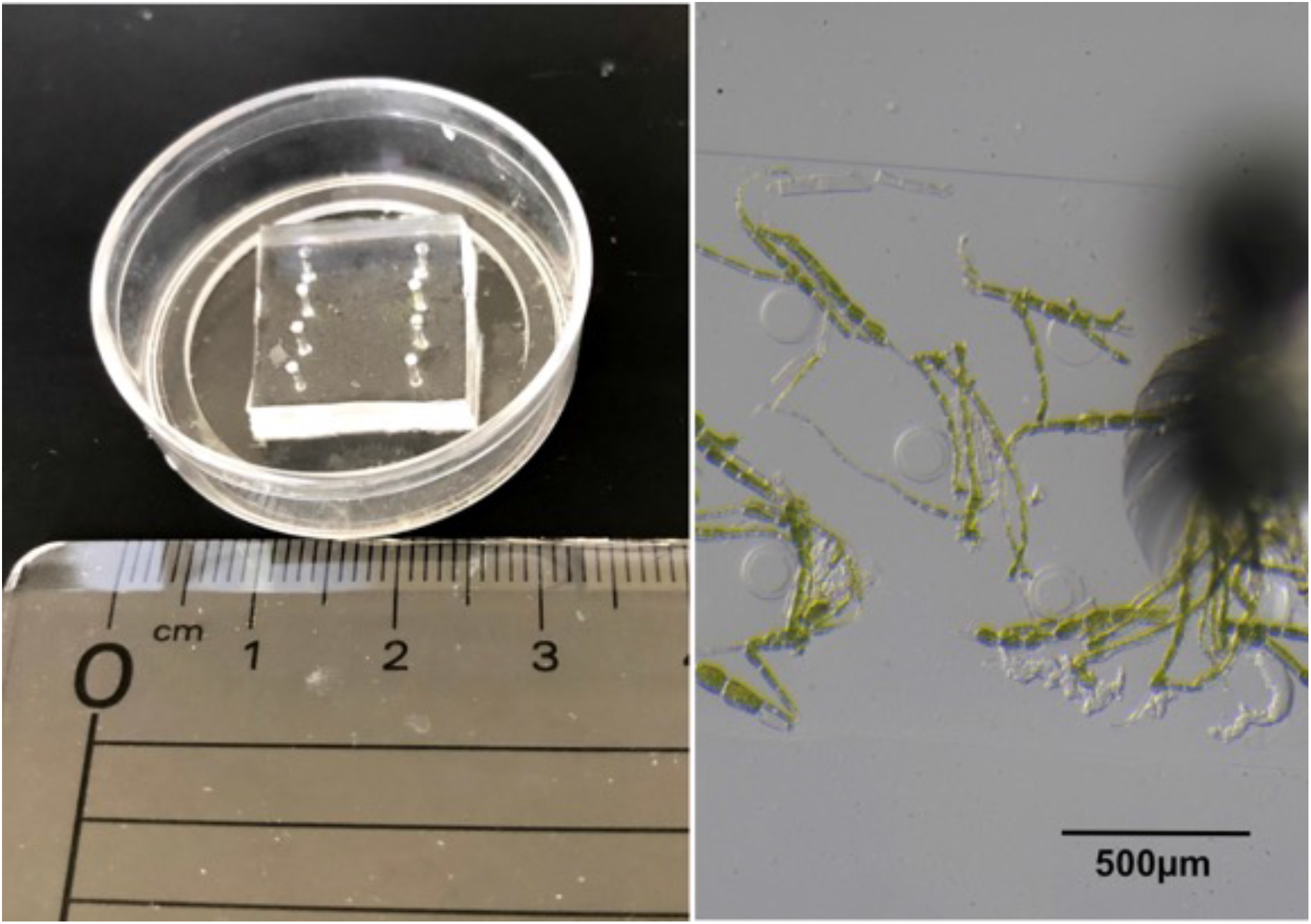
Microfluidic device for live-cell imaging of macroalgae. (Left) A PDMS device, 15 µm in height, was attached to the glass-bottom dish. (Right) Severed thalli of *Cladosphora* were injected into the device.

**Figure S2.**
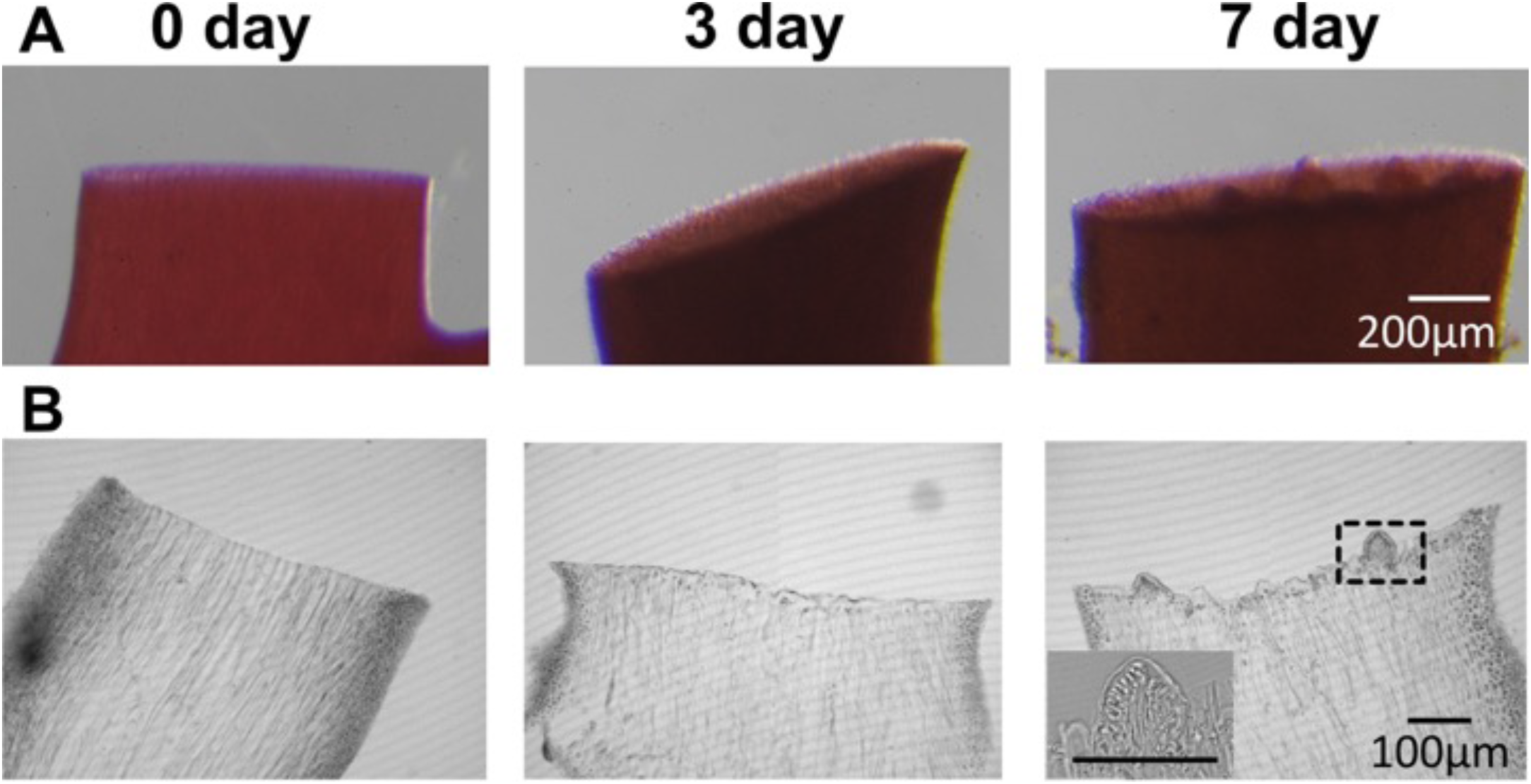
Budding at the severed surface in *Gelidium elegans*. (A) Stereomicroscopic images of the severed surface of *G. elegans* thalli at day 0 (immediately after severing), 3, and 7. (B) Frozen section images of a cut surface of *G. chorda* thalli. Inset: enlarged image.

## Notes

### Competing Interest Statement

The authors have declared no competing interest.

